# MicroRNA inhibition using antimiRs in acute human brain tissue sections

**DOI:** 10.1101/2022.04.05.487136

**Authors:** Gareth Morris, Elena Langa, Conor Fearon, Karen Conboy, Kelvin Lau E-How, Amaya Sanz-Rodriguez, Donncha F O’Brien, Kieron Sweeney, Austin Lacey, Norman Delanty, Alan Beausang, Francesca M Brett, Jane B Cryan, Mark O Cunningham, David C Henshall

## Abstract

**Introduction:** An emerging pre-clinical approach for the treatment of pharmacoresistant epilepsy is targeting the microRNA (miRNA) system. MiRNAs are short noncoding RNAs that suppress gene expression at the post-transcriptional level. Targeting miRNAs, which is possible using antisense oligonucleotide ‘antimiRs’ can produce broad effects on gene expression suited to the complex pathophysiology in temporal lobe epilepsy. Potent anti-seizure and disease-modifying effects have been reported for antimiRs targeting microRNA-134 (antimiR-134). To date, however, pre-clinical testing has been performed using *in vitro* cell cultures and rodent models. It is uncertain how well this approach will translate to the clinic. Here, we develop an antimiR testing platform in human brain tissue sections.

**Methodology:** Human brain specimens were obtained with consent from patients undergoing resective surgery to treat focal drug-resistant epilepsy. Neocortical specimens were submerged in modified artificial cerebrospinal fluid (ACSF), dissected for clinical neuropathological examination, and unused material transferred for sectioning. Individual tissue sections were incubated in oxygenated ACSF, containing either antimiR-134 or a non-targeting control antimiR, for 24 hours at room temperature. RNA integrity was assessed using BioAnalyzer processing, and individual miRNA levels measured using RT-qPCR.

**Results:** ACSF transport had no obvious impact on any clinical neurosurgical or neuropathological procedure and specimens were confirmed to be viable following this process. RNA was well-preserved by transportation of specimens in ACSF, with RNA integrity scores significantly higher than tissue transported without ACSF. AntimiR-134 mediated a specific and dose- dependent knockdown of miR-134 in human neocortical sections, with approximately 75% reduction of miR-134 at 1 µM and 90% reduction at 3 µM. These doses did not have off-target effects on expression of a selection of three other miRNAs (miR-10, miR-129 or miR- 132).

**Significance:** This is the first demonstration of antimiR-134 effects in live human brain tissues. The findings lend further support to the preclinical development of miR-134 and offer a flexible platform for the pre-clinical testing of antimiRs, and other antisense oligonucleotide therapeutics, in human brain.

**Key points:**

- ASO antimiRs are promising treatments for pharmacoresistant epilepsy
- We developed a pipeline to preserve live human neocortical brain specimens from people undergoing resective surgery
- RNA integrity was sufficient to measure miRNA levels in human brain tissues transported in modified ACSF
- Incubation of acute human neocortical specimens in antimiR-134 resulted in potent and specific reduction in miR-134 levels
- Acute human brain slices are a promising model for testing ASOs

## Introduction

Epilepsy is a chronic neurological disease which affects an estimated 70 million people worldwide^1^. Epilepsy manifests clinically as seizures, alongside potential cognitive and psychiatric comorbidities^2^. The frontline treatment for epilepsy is with anti-seizure medications (ASMs), of which around 30 are clinically available^3,4^. Despite the range of ASMs available, 30% of patients continue to experience seizures despite optimal treatment profiles - these cases are classified as pharmacoresistant^5^. Uncontrolled seizures are a critical risk factor for sudden unexpected death in epilepsy (SUDEP)^6^. Moreover, ASMs must be taken daily, do not address disease co-morbidities, and are sometimes associated with significant adverse effects^7^. Therefore, there is an urgent and unmet requirement to develop novel therapeutic strategies in epilepsy which meet clinical and patient needs.

One leading preclinical solution to this problem targets the microRNA (miRNA) system^8–11^. MiRNAs are endogenous ∼22nt noncoding RNAs which repress the translation of targeted mRNAs via complementary binding to targets regions in the 3’ UTR^12^. MiRNA dysregulation is linked to epilepsy^8–11,13^ and individual miRNAs may represent either biomarkers or therapeutic anti-seizure targets. Antisense oligonucleotides (ASO) targeting miRNAs, termed ‘antimiRs’ provide potent, specific and lasting in vivo knockdown of their targets. Among leading approaches in epilepsy, inhibition of miR-134 with antimiRs has been reported to produce potent anti-seizure and disease-modifying effects in rodent models of seizures and epilepsy^14–18^. However, it remains unclear whether effects of antimiR-134 will translate to the clinic. Notably, while the sequence of miR-134 itself is conserved between rodent and human, some of its target mRNAs are not^19^. It is therefore unknown whether antimiR-134 will have the same molecular, biophysical and anti-seizure effects in human brain. Further, it is unclear how efficiently antimiR-134 will penetrate into human brain cells. ASO uptake mechanisms can vary between cell types^20^ and may not be the same in rodent and human brain.

A platform to test this is offered by resective brain surgeries for pharmacoresistant epilepsy^21,22^. Surgical removal of the epileptogenic zone (the area of cortex that is indispensable for the generation of epileptic seizures^23^) can be indicated in pharmacoresistant focal epilepsies in which the epileptogenic zone does not overlap with eloquent cortex^24^. The resected brain tissue can be collected and used for research purposes, offering clear scientific advantages over rodent epilepsy models, most notably superior construct validity due to the use of human brain to study a human condition^21^. Resected human brain tissue can be processed acutely for up to 72 hours^25^, or maintained for longer periods of time in organotypic slice culture^26,27^. This provides an ideal preparation to test the molecular effects of antimiR-134 in the human brain.

Here, we set out an adapted methodology to transport human brain tissue from the neurosurgery theatre to the research laboratory for molecular assessment, without impacting key clinical processes or diagnosis. We verified that specimens processed in this way were viable and had sufficient RNA integrity for molecular studies. We used sections of these specimens to inhibit targeted miRNA in human brain using antimiR-134 as a proof of concept. This provides new tools for molecular and specifically miRNA research in the epileptic human brain.

## Methods

### Ethical approvals

All studies using human brain tissue were approved by the local ethical committee at Beaumont Hospital Dublin (Reference Number 13/75, 20/58). All patients approached for this study had the capacity to give fully informed consent and were provided with a Patient Information Leaflet to read before providing consent. Tissues used were removed as part of the normal clinical procedure and no extra tissue is ever resected for research use.

### Resective neurosurgery

All patients enrolled in the study underwent resective brain surgery at Beaumont Hospital, Dublin, to alleviate focal epilepsy. Patient and specimen details can be found in Table 1. Neocortical tissues were resected *en bloc* as part of the surgical procedure. We immediately submerged these specimens into ice cold oxygenated sucrose artificial cerebrospinal fluid (ACSF; in mM: 205 sucrose, 10 glucose, 2.5 KCl, 5 MgCl_2_, 0.1 CaCl_2_, NaHCO_3_, NaH_2_PO_4_.H_2_O) and transported them in sucrose ACSF and on ice to the neuropathology laboratory as part of the clinical process. This transport process took approximately five minutes, during which time the ACSF was not oxygenated further.

**Table 1:** Details of patients and tissue specimens used

| **Patient Research ID** | **Age (years)** | **Sex** | **Nature of initial specimen** |
| --- | --- | --- | --- |
| **1** | 37 | M | LTL |
| **2** | 61 | F | LTL |
| **3** | 59 | M | LTL |
| **4** | 62 | M | Tumour |
| **5** | 60 | M | LTL |
| **6** | 43 | M | LTL |
| **7** | 39 | F | FL |
| **8** | 22 | F | LTL |
| **9** | 54 | M | LTL |
| **10** | 47 | F | LTL |
| **11** | 39 | M | LTL |
| **12** | 59 | M | LTL |
| **13** | 28 | M | LTL |

**Legend:** LTL - lateral temporal lobe; FL - frontal lobe

### Neuropathological assessment

Neocortical specimens were removed from the ACSF and assessed for diagnosis. After macroscopic examination and serial sectioning, a small piece of the specimen (approx. 1×1×0.5 cm) was immediately re-submerged in ACSF and transported on ice to the laboratory for research purposes (a further 2-3 minutes, without further oxygenation of the ACSF). A distinction was made between peripheral sections in immediate contact with ACSF, and deeper/ mid sections, largely remote from ACSF. The remaining specimen was fixed overnight in 10% formalin after which they were processed and embedded in paraffin. Sections (4 µm thick) were stained with haematoxylin and eosin (H&E) +/- GFAP, MAP2, NeuN immunohistochemistry for systematic assessment of neuronal loss, astrogliosis, malformations of cortical development (e.g. cortical neuronal dyslamination, dysmorphic neurons, balloon cells and oligodendroglial hyperplasia) as well as vascular malformations, hamartomas or neoplastic processes and ischaemic or inflammatory lesions. We compared the quality and robustness of neuropathological assessment in peripheral sections that were in direct contact with ACSF against the deeper sections, otherwise processed in the same way.

### AntimiR treatment of human brain specimens

On arrival in the research laboratory, the sucrose ACSF containing the specimen was immediately re-oxygenated. The specimen was first transferred to a petri dish filled with fresh ice cold and oxygenated sucrose ACSF. Care was taken to remove any remaining *pia mater* from the specimen. For molecular assessments, the specimen was divided into cubes (approx. 1 mm^3^) using a scalpel. We created a low volume incubation chamber for these specimens using a standard 12 well plate with small mesh inserts. Specimens were placed into the mesh inserts, which were then submerged into 4 mL normal ACSF (in mM: 125 NaCl, 10 Glucose, 3 KCl, 2 CaCl_2_, 1 MgCl_2_, 26 NaHCO_3_, 1.25 NaH_2_PO_4_.H_2_O) within one well of the plate. The low volume used facilitated the application of physiologically relevant miRNA inhibitors. Specimens were incubated for 24 hours at room temperature in the presence of either an ASO antimiR targeting hsa-miR-134-5p (‘Ant-134’; Qiagen, Manchester, UK; catalogue number 339132: TGGTCAACCAGTCACA), or a scrambled (Scr) nontargeting control antimiR (‘Negative control A’; Qiagen; catalogue number 339136: TAACACGTCTATACGCCCA), at 0.1, 1 and 3 µM. After 24 hours specimens were removed from ACSF and flash frozen at -80 °C.

### Rodent brain specimens

As part of another study, C57BL/6 mice (age range p21-p48) underwent unilateral stereotaxic injection of PBS into the amygdala. 24 hours later, mice were euthanised with 0.2 ml IP sodium pentobarbital (200 mg/ml). Mice were then perfused with 20 ml PBS via cardiac puncture, and brain tissue frozen at -80 °C for RNA extraction.

### RNA extraction

Tissue was homogenized in 800 µL of Trizol (Qiagen, UK) and centrifuged at 12,000 g for 10 min at 4°C. 200 µl of chloroform (Sigma-Aldrich, Ireland) was added to each sample, which were vigorously mixed for 15 seconds and incubated at room temperature for 3 minutes. Samples were then centrifuged at 13,000 rpm for 15 min at 4 °C. The upper aqueous phase was transferred to a new tube and 450 µL of isopropanol (Sigma-Aldrich, Ireland) was added. Samples were incubated at − 20°C overnight and then centrifuged at 13,000 rpm for 15 min at 4°C. The supernatant was removed and the RNA pellets were washed twice with 750 µL of 75% cold ethanol, centrifuged at 12,000 g for 5 min and the ethanol removed. Pellets were allowed to dry for 1 h and resuspended in 20 µL of RNase free water. Samples were incubated for 10 min at 60°C, shaking at 800 rpm, and stored at - 80°C.

### RNA integrity

RNA integrity and quality was assessed using the Agilent 2100 Bioanalyzer instrument (Agilent) using the RNA 6000 Nano kit (5067-1511) according to manufacturers guidelines. In short, reagents were equilibrated to room temperature followed by preparation of the gel-dye mix and ladder. The microfluidic RNA Nano chip was prepared followed by loading of 1 µl of sample. Total RNA quality was assessed by determining the ratio of ribosomal peaks (18S/28S). This is used to derive the RNA Integrity Number (RIN) value, which provides an objective readout of RNA quality between 0 (completely degraded) and 10 (highly intact RNA).

### MiRNA expression

250 ng of total RNA was reverse transcribed using stem-loop Multiplex primer pools (Applied Biosystems, Dublin, Ireland). We used reverse-transcriptase-specific primers for mmu-miR-134 (Applied Biosystems miRNA assay ID 001186; also targets hsa-miR-134-5p), hsa-miR-10 (Applied Biosystems miRNA assay ID 000387), hsa-miR-129 (Applied Biosystems miRNA assay ID 000590) and hsa-miR-132 (Applied Biosystems miRNA assay ID 000457).

Real-time quantitative PCR was carried out on a QuantStudio™ 12K Flex PCR system (Applied Biosystems) using TaqMan miRNA assays (Applied Biosystems). RNU19 (Applied Biosystems miRNA assay ID 001003) was used for normalisation. A relative fold change in expression of the target gene transcript was determined using the comparative cycle threshold method (2−ΔΔCT).

### Statistics

Data were tested for normality using a Kolmogorov-Smirnov test. All quantitative comparisons made contained at least one non-normally distributed dataset, and so all data were analysed using a Kruskal-Wallis test with Dunn’s testing for multiple comparisons. Averages are shown as median ± interquartile range (IQR). Statistical analyses were performed using GraphPad Prism v9.3.1 (GraphPad Software, CA, USA). Individual samples were randomised to experimental groups. RNA extraction and RT-qPCR were performed blind to experimental group.

## Results

### Tissue specimens obtained

A total of 13 human neocortical brain specimens were obtained from patients undergoing resective surgery for epilepsy. This included nine male and four female patients, with an age range from 22-62 years. See Table 1 for details of individual specimens used. Specimens were obtained by the researcher directly in the surgical theatre, with the researchers trained to safely enter the theatre with no impact on the clinical procedure (overview in Figure 1).

**Figure 1:**
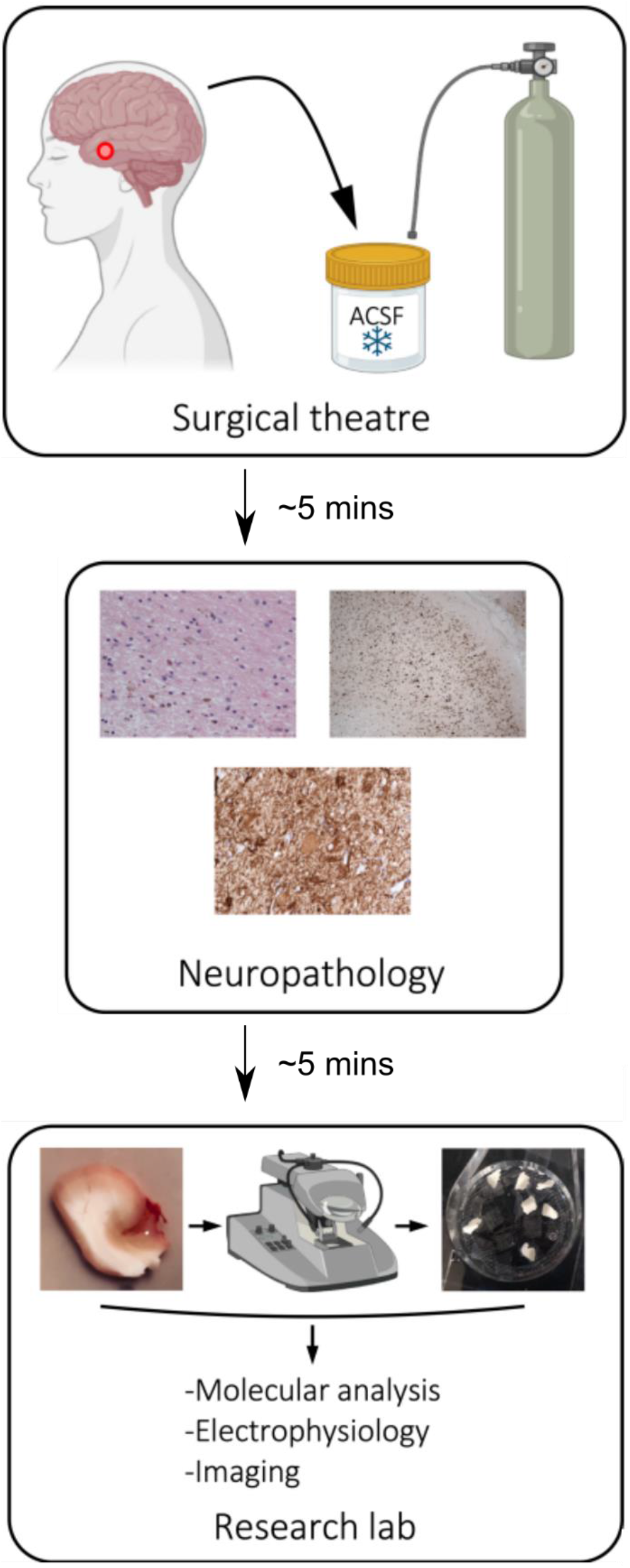
Schematic overview of brain tissue processing from neurosurgery to research lab. Resected human brain tissue specimens were obtained directly from the neurosurgical theatre and immediately submerged into oxygenated ice cold ACSF. Specimens were transported in ACSF to the neuropathology laboratory as part of the clinical pathway. A proportion of the specimen was then returned to ACSF and taken to the research lab for study.

### Neuropathological assessment

We first verified that transporting surgical brain specimens in ACSF did not impact key neuropathological assessments and diagnosis as part of the clinical pathway. Qualitative analysis of sections transported in ACSF showed conserved cellular morphology as observed with both H&E and immunohistochemical staining protocols. A number of different pathologies could be identified from specimens transported in this way, including perioperative dark cell change (H&E, Figure 2A), subpial gliosis with upregulation of GFAP (Figure 2B); oligodendroglial hyperplasia (Figure 2C); focal cortical dysplasia type 2B (Figures 2D-F) with abnormal cortical neuronal lamination (Figure 2D,E) and dysmorphic neurons and balloon cells (Figure 2F); remote cortical haemorrhage and gliosis (Figure 2G) with white matter cavitation (Figure 2H). Figure 2I shows a comparable section with remote haemorrhage and gliosis obtained from a specimen transported without direct ACSF contact, showing no qualitative difference caused by our ACSF transport approach.

**Figure 2:**
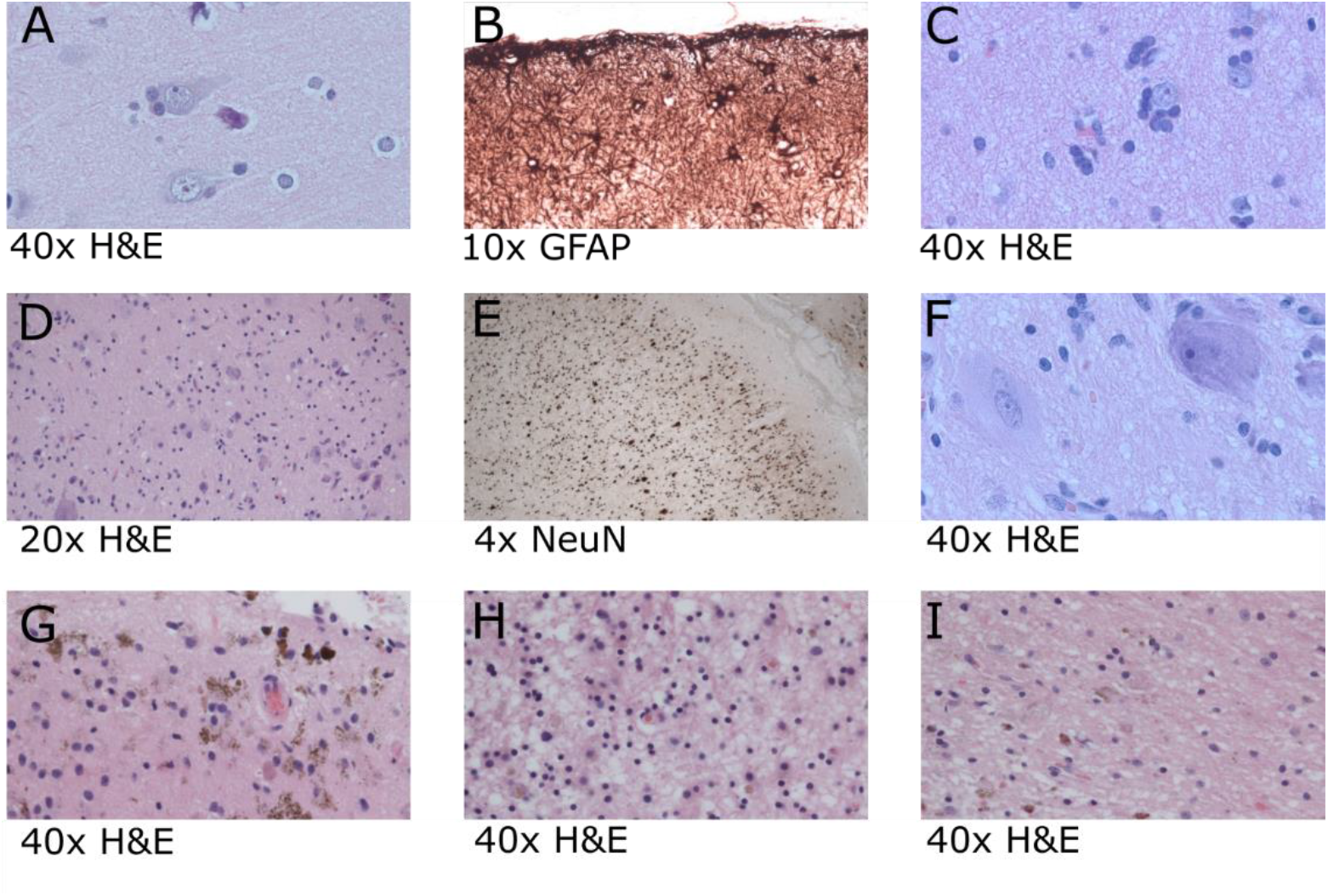
Sucrose ACSF transport procedures have no impact on neuropathological assessments. Examples of neuropathological observations made in specimens transported in ACSF illustrate the compatibility of our approach with the clinical pathway. (A) normal cortex and neuronal morphology with perioperative ‘dark cell change’ (B) –subpial gliosis; (C) – oligodendroglial hyperplasia; (D,E,F) – focal cortical dysplasia type 2B characterised by disorganised lamination (D,E H&E & NeuN) and dysmorphic neurons and balloon cells (F); (G,H) – remote haemorrhage and gliosis. (I) shows a directly comparable section exhibiting remote haemorrhage and gliosis in a specimen handled in a standard way without direct ACSF exposure, indicating no qualitative impact of ACSF transport.

### Tissue viability and RNA Integrity

We then sought to verify that these specimens were viable and suitable for use as a pre-clinical research model for antimiR testing. We first assessed the RNA integrity in a subset of samples transported in ACSF, compared with equivalent neocortical samples brought to the lab without being submerged in any solution, using previous methods in the group. BioAnalyzer processing revealed significantly higher RNA Integrity Number (RIN) values in samples processed using ACSF (Figure 3). Notably, these RIN values were not significantly different from those obtained in rodent tissues processed for molecular biology. This provides an objective quality control measure validating the use of our approach as a platform to study RNA-targeting oligonucleotide therapeutics. However, it must also be considered that the RNA yield from human brain tissue acquired in this way (Figure 3B) was lower than in ‘gold-standard’ rodent brain tissue prepared for molecular biology (Figure 3C).

**Figure 3:**
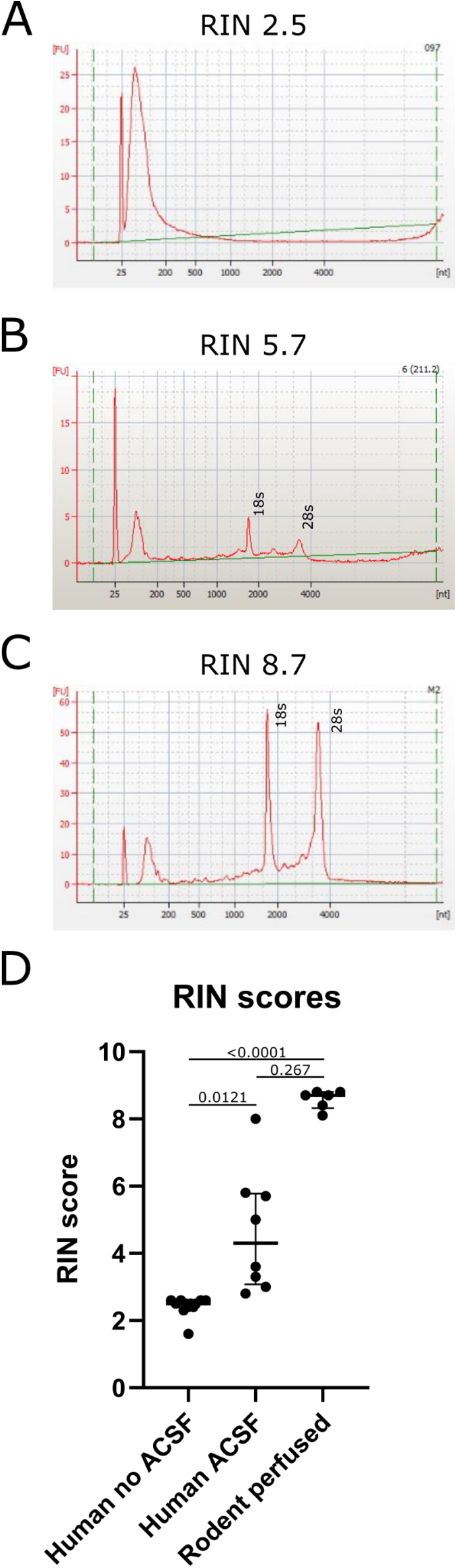
Human neocortical specimens transported in sucrose ACSF are viable for molecular experiments. BioAnalyzer traces from human brain tissue transported without (A) ACSF and with (B) ACSF. (C) A trace from a mouse brain perfused for molecular biology is included as a ‘gold standard’ comparison. Note that the RNA yield in (B) is much lower than in (C). (D) RNA Integrity Number values derived from Bioanalyzer processing provide an objective measure of RNA quality. Human samples processed using our method had significantly higher RIN values than those processed using standard methods (Kruskal-Wallis test with Dunn’s multiple comparisons test).

### MicroRNA-134 inhibition in human brain tissue

Having confirmed the viability and RNA integrity in our specimens, we next asked whether ant-134 would efficiently transfect human brain cells and inhibit miR-134. We applied ant-134 for 24 hours (Figure 4A,B) at a range of concentrations which were based on our previous findings in induced pluripotent stem cell (iPSC)-derived neurons. Using RT-qPCR, we observed a dose-dependent inhibition of miR-134, with significant knockdown mediated by 1 µM and 3 µM ant-134, compared with non-targeting control antimiR (Figure 4C). Finally, we verified that these viable concentrations of ant-134 did not cause off-target effects on examples of other brain-enriched miRNAs. RT-qPCR analysis of the same samples showed that the expression of miR-10, miR-129 and miR-132 were all unaffected by ant-134 (Figure 4D). Together, this indicated that our approach can provide a platform for the testing of antimiR uptake and efficacy in resected human brain tissue, offering an important tool in pre-clinical research to aid in the translation of oligonucleotide therapeutics to the clinic.

**Figure 4:**
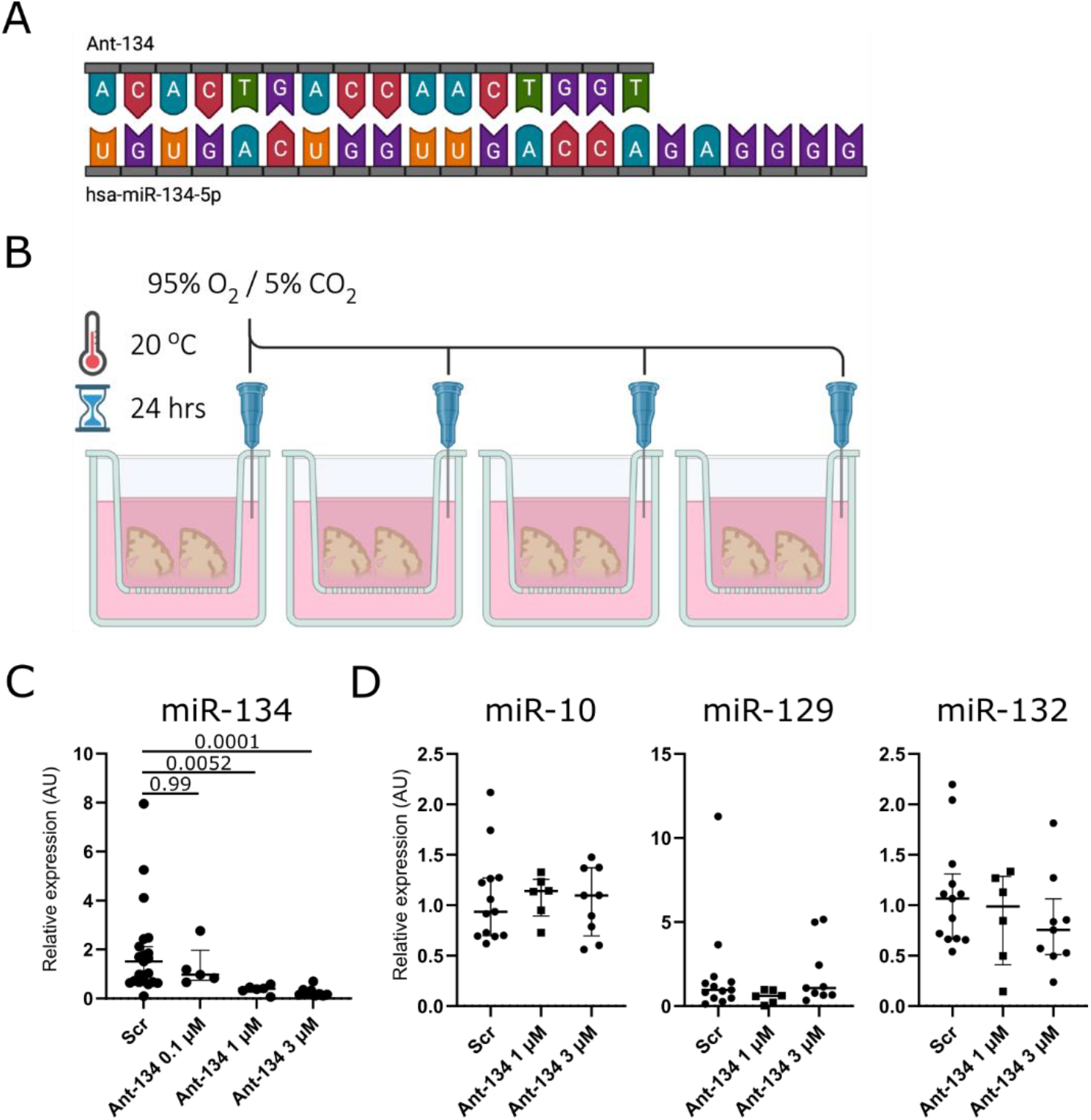
Ant-134 mediates a dose-dependent knockdown of microRNA-134 in human neocortex. (A) Sequences of hsa-miR-134-5p (22-mer) and ant-134 (16-mer) indicates perfect complementarity between the two. (B) Experimental set-up to treat acutely sectioned human brain specimens with antimiR. Sections were placed into small inserts with a permeable mesh at the bottom. Inserts were placed into individual wells of a standard 12- well plate and submerged into 4 mL normal ACSF containing ant-134 or scr at varying concentrations. The ACSF is each well was oxygenated using a syringe needle connected to a carbogen gas supply. This preparation was left for 24 hours at room temperature. (C) RT-qPCR shows robust dose-dependent knockdown of miR-134 after 24 hours (Kruskal-Wallis test with Dunn’s test for multiple comparisons). (D) For the viable doses of ant-134, we did not observe off-target inhibition of either miR-10, miR-129 or miR-132 (all Kruskal-Wallis test with Dunn’s multiple comparisons tests)..

## Discussion

Whilst miRNA targeting using antimiRs is a promising pre-clinical strategy to treating pharmacoresistant epilepsy, testing to date has focused on rodent models. This has the drawback that antimiR uptake and efficacy may be different in human brain tissue. Here, we demonstrate for the first time that antimiRs can mediate specific and dose-dependent target knockdown in human neocortical specimens.

### AntimiR testing in human brain - advantages over other epilepsy models

The approach outlined provides a new methodology to interrogate the molecular effects of miRNA manipulation in the human brain. This provides a much-needed tool to aid translation of miRNA-targeting therapies to the clinic^8,21^. The primary advantage of this method is that it ‘models’ epilepsy in tissues containing human, rather than rodent genetics. This is particularly applicable to studies using miRNA-based approaches because the sequences of miRNAs and their mRNA targets are not always conserved. For example in the case of miR-134, the interaction between miR-134 and *Limk1*, thought to be responsible in part for its anti-seizure effect^14,17^, may not be conserved^19^. Therefore, studies on human-derived tissues are key to test this. Further, molecular testing in human brain tissues is an important regulatory step. Because miRNA:mRNA interactions are not always conserved, there is the possibility that miRNA manipulation in humans could give rise to unanticipated adverse effects, not observed in rodents, due to the greater complexity of the human brain. Further, though beyond the scope of the present study, human brain slices provide a tool to induce epileptiform activity acutely for studies into mechanisms and treatment of seizures^21,22,25,27,28^. This presents the opportunity to test novel therapies in real diseased tissue, rather than in rodent proxies of epilepsy. Alternative human-based approaches to epilepsy research use iPSCs^29–35^. These are human-derived precursors which can be differentiated into mature neurons in 2D culture, or grown into 3D brain organoids^36,37^. However, questions remain about the physiological relevance of such models, as they do not form physiologically realistic circuits and so cannot capture the full pathophysiology of epilepsy^38^. Therefore, real human brain tissue, obtained from patients undergoing surgery, remains the ultimate tool in translational epilepsy research^21^. Finally, there are also ethical advantages to our approach, which represents a replacement for animal research within the concept of the 3Rs^39^.

### A human brain-based platform for ASO translation

Whilst the present study focused on antimiR technology, our approach could be applied more broadly to other ASO strategies. Due to their relative ease of delivery and ability to readily target widespread brain structures, ASOs are emerging as a key approach to treating epilepsies – particularly those arising from gene mutations that thus effect the whole brain – and other neurological diseases. For example, ASO gapmers, which typically favour target mRNA degradation, have been used in a mouse model of a genetic channelopathy which leads to epilepsy^40^. Current challenges for ASO translation from animal models to human clinical applications include on- and off-target toxicity effects^41^, which may be unique to the human brain and not possible to observe in mice. ASO mechanisms of action^41^ can include steric blocking of mRNA translation, degradation of target mRNA or regulation of RNA splice events^42^. Different ASOs may also have different lengths and chemistries, which can affect their uptake, bioavailability and cellular effects^8^. Our methodology provides a platform which could be used to test all of these ASO mechanisms and properties in real human brain tissue. Further, our approach can be applied to other neurological diseases beyond epilepsy. The neurosurgical procedure typically involves the resection of non-epileptic temporal neocortex, in order to access the deeper epileptogenic zone. Therefore, this tissue is largely representative of healthy human neocortex, and so could be used to probe the molecular mechanisms of ASOs for other diseases, where their mechanism might be impacted by the diseased nature of the epileptogenic zone. ASOs have shown great promise in diseases including spinal muscular atrophy^43,44^ and Huntington’s disease^45^, although clinical trials for the latter have recently been halted^46^. We propose that resected human brain tissue could be used as an additional layer of pre-clinical testing in ASO discovery pipelines, to aid in the interrogation of possible adverse effects that cannot be predicted in other models.

### Integration into the neurosurgical and neuropathological workflow

Our strategy to collect human brain tissue from resective epilepsy surgeries fitted seamlessly into the clinical pathway as a result of strong collaboration between neurologists, neurosurgeons, neuropathologists, medical laboratory scientists and researchers. Our researchers were trained to safely enter the surgical theatre to collect tissue, meaning that the approach did not place any additional demands on the surgical team. This had the added advantage that resected specimens were immediately transferred to fresh sucrose ACSF and could be moved by the researcher to the neuropathology laboratory and the research laboratory as quickly as possible, maximising tissue viability. We also worked closely as a multidisciplinary team to verify that sucrose ACSF transportation had no impact on neuropathological assessments. Together, the training of our researchers to work in the clinical environment, alongside close collaboration with our neurosurgeons and neuropathologists, was critical to our approach.

### Limitations of the approach

Our approach, and the use of human tissue in epilepsy research in general, also has limitations. Notably, the resected tissues are also required for neuropathologic diagnosis. Whilst we present a methodology to fully integrate our approach into clinical pathways, it remains sub-optimal from a research viewpoint that tissues cannot be taken directly to the research lab. This could have some impact on tissue viability for research, though our assessments suggest that human tissues, when handled carefully, likely have comparable quality to freshly prepared rodent samples. Another challenge is the relatively limited availability and the heterogeneity of the specimens available. Our approach relies upon a research laboratory in close physical proximity to a specialist neurosurgery centre. The frequency of surgeries dictates tissue availability and may not always be consistent due to the nature of the clinical pathway. Coupled with this, specimens are highly heterogeneous and are obtained from patients with different sex, age and ASM histories. This likely adds more variability to datasets and necessitates higher sample sizes, although this is mitigated in part because it is usually possible to obtain a large number of acute sections from one individual resected specimen. Taken together, the relatively low throughput and need for more samples means human tissue-based studies take much longer than those in rodents, and this must be considered by researchers before beginning such studies. Finally, from the point of view of modelling seizures acutely in human tissue slices, these specimens are studied in isolation. This has the disadvantage that they are dissociated from wider brain networks, prohibiting the study of wider seizure propagation pathways, and that it is not possible to study peripheral effects of experimental therapies.

## Conclusions

We present a method by which we incorporated the use of resected human brain tissues for research into pre-existing clinical pathways. These tissues are amenable to studies including molecular and electrophysiological assessment, and provide a critical tool in translational epilepsy research. Further, we demonstrated for the first time the use of ant-134, a leading preclinical candidate for pharmacoresistant epilepsy, in human brain. This paves the way for further study into the mechanisms of ant-134, and other miRNA-based approaches, in human brain.

## Acknowledgments

This publication has emanated from research supported in part by a research grant from Science Foundation Ireland (SFI) under Grant Number 16/RC/3948 and co-funded under the European Regional Development Fund and by FutureNeuro industry partners. GM is supported by a Marie Skłodowska-Curie Actions Individual Fellowship (‘EpimiRTherapy’, H2020-MSCA-IF-2018 840262) and an Emerging Leader Fellowship Award from Epilepsy Research UK (grant reference F2102 Morris).

## Reference list

1. Ngugi AK, Bottomley C, Kleinschmidt I, Sander JW, Newton CR. Estimation of the burden of active and life-time epilepsy: A meta-analytic approach. Epilepsia. 2010; 51(5):883–90.

2. Mula M, Kanner AM, Jetté N, Sander JW. Psychiatric Comorbidities in People With Epilepsy. Neurol Clin Pract [Internet]. 2021; 11(2):e112–20. Available from: http://www.ncbi.nlm.nih.gov/pubmed/33842079

3. Löscher W. Single-Target Versus Multi-Target Drugs Versus Combinations of Drugs With Multiple Targets: Preclinical and Clinical Evidence for the Treatment or Prevention of Epilepsy. Frontiers in Pharmacology. 2021; 12(October):1–22.

4. Chen Z, Brodie MJ, Liew D, Kwan P. Treatment outcomes in patients with newly diagnosed epilepsy treated with established and new antiepileptic drugs a 30-year longitudinal cohort study. JAMA Neurology. 2018; 75(3):279–86.

5. Schmidt D, Löscher W. Drug resistance in epilepsy: Putative neurobiologic and clinical mechanisms. Epilepsia. 2005; 46(6):858–77.

6. Hesdorffer DC, Tomson T, Benn E, Sander JW, Nilsson L, Langan Y, et al. Combined analysis of risk factors for SUDEP. Epilepsia. 2011; 52(6):1150–9.

7. Perucca P, Gilliam FG. Adverse effects of antiepileptic drugs. The Lancet Neurology [Internet]. 2012; 11(9):792–802. Available from: https://linkinghub.elsevier.com/retrieve/pii/S1474442212701539

8. Morris G, O’Brien D, Henshall DC. Opportunities and challenges for microRNA-targeting therapeutics for epilepsy. Trends in Pharmacological Sciences [Internet]. 2021; 42(7):605–16. Available from: https://doi.org/10.1016/j.tips.2021.04.007

9. Henshall DC. MicroRNA and epilepsy: Profiling, functions and potential clinical applications. Current Opinion in Neurology. 2014; 27(2):199–205.

10. Brennan GP, Henshall DC. MicroRNAs as regulators of brain function and targets for treatment of epilepsy. Nat Rev Neurol [Internet]. 2020; 16(9):506–19. Available from: http://www.ncbi.nlm.nih.gov/pubmed/32546757

11. Henshall DC, Hamer HM, Pasterkamp RJ, Goldstein DB, Kjems J, Prehn JHM, et al. MicroRNAs in epilepsy: pathophysiology and clinical utility. The Lancet Neurology [Internet]. 2016; 15(13):1368–76. Available from: http://dx.doi.org/10.1016/S1474-4422(16)30246-0

12. Bartel DP. Metazoan MicroRNAs. Cell. 2018; 173(1):20–51.

13. Henshall DC. MicroRNAs in the pathophysiology and treatment of status epilepticus. Front Mol Neurosci. 2013; 6(November).

14. Jimenez-Mateos EM, Engel T, Merino-Serrais P, McKiernan RC, Tanaka K, Mouri G, et al. Silencing microRNA-134 produces neuroprotective and prolonged seizure-suppressive effects. Nat Med [Internet]. 2012; 18(7):1087–94. Available from: http://www.pubmedcentral.nih.gov/articlerender.fcgi?artid=3438344&tool=pmcentrez&rendertype=abstract

15. Jimenez-Mateos EM, Engel T, Merino-Serrais P, Fernaud-Espinosa I, Rodriguez-Alvarez N, Reynolds J, et al. Antagomirs targeting microRNA-134 increase hippocampal pyramidal neuron spine volume in vivo and protect against pilocarpine-induced status epilepticus. Brain Structure and Function. 2015; 220(4):2387–99.

16. Reschke CR, Silva LFA, Norwood BA, Senthilkumar K, Morris G, Sanz-Rodriguez A, et al. Potent Anti-seizure Effects of Locked Nucleic Acid Antagomirs Targeting miR-134 in Multiple Mouse and Rat Models of Epilepsy. Molecular Therapy - Nucleic Acids. 2017; 6:45–56.

17. Morris G, Reschke CR, Henshall DC. Targeting microRNA-134 for seizure control and disease modification in epilepsy. EBioMedicine [Internet]. 2019; 45:1–9. Available from: https://doi.org/10.1016/j.ebiom.2019.07.008

18. Morris G, Brennan GP, Reschke CR, Henshall DC, Schorge S. Spared CA1 pyramidal neuron function and hippocampal performance following antisense knockdown of microRNA-134. Epilepsia. 2018; (May):1–9.

19. Schratt GM, Tuebing F, Nigh EA, Kane CG, Sabatini ME, Kiebler M, et al. A brain-specific microRNA regulates dendritic spine development. Nature. 2006; 439(7074):283–9.

20. Crooke ST, Wang S, Vickers TA, Shen W, Liang XH. Cellular uptake and trafficking of antisense oligonucleotides. Nature Biotechnology. 2017; 35(3):230–7.

21. Morris G, Rowell R, Cunningham MO. Limitations of animal epilepsy research models: Can epileptic human tissue provide translational benefit? ALTEX [Internet]. 2021;. Available from: https://www.altex.org/index.php/altex/article/view/2003

22. Jones RSG, da Silva AB, Whittaker RG, Woodhall GL, Cunningham MO. Human brain slices for epilepsy research: Pitfalls, solutions and future challenges. Journal of Neuroscience Methods [Internet]. 2016; 260:221–32. Available from: http://dx.doi.org/10.1016/j.jneumeth.2015.09.021

23. Rosenow F, Lüders H. Presurgical evaluation of epilepsy. Vol. 124, Brain. 2001.

24. Jobst BC, Cascino GD. Resective epilepsy surgery for drug-resistant focal epilepsy: A review. JAMA - Journal of the American Medical Association. 2015; 313(3):285–93.

25. Wickham J, Brödjegård NG, Vighagen R, Pinborg LH, Bengzon J, Woldbye DPD, et al. Prolonged life of human acute hippocampal slices from temporal lobe epilepsy surgery. Scientific Reports. 2018; 8(1):1–13.

26. Schwarz N, Uysal B, Welzer M, Bahr JC, Layer N, Löffler H, et al. Long-term adult human brain slice cultures as a model system to study human CNS circuitry and disease. Elife. 2019; 8:1–26.

27. Wickham J, Corna A, Schwarz N, Uysal B, Layer N, Honegger JB, et al. Human Cerebrospinal Fluid Induces Neuronal Excitability Changes in Resected Human Neocortical and Hippocampal Brain Slices. Frontiers in Neuroscience. 2020; 14(April):1–14.

28. Gabriel S, Njunting M, Pomper JK, Merschhemke M, Sanabria ERG, Eilers A, et al. Stimulus and potassium-induced epileptiform activity in the human dentate gyrus from patients with and without hippocampal sclerosis. Journal of Neuroscience. 2004; 24(46):10416–30.

29. Engle SJ, Blaha L, Kleiman RJ. Best Practices for Translational Disease Modeling Using Human iPSC-Derived Neurons. Neuron [Internet]. 2018; 100(4):783–97. Available from: https://doi.org/10.1016/j.neuron.2018.10.033

30. Dolmetsch R, Geschwind DH. The human brain in a dish: The promise of iPSC-derived neurons. Cell [Internet]. 2011; 145(6):831–4. Available from: http://dx.doi.org/10.1016/j.cell.2011.05.034

31. Bardy C, Hurk M van den, Kakaradov B, Erwin JA, Jaeger BN, Hernandez R v, et al. Predicting the functional states of human iPSC-derived neurons with single-cell RNA-seq and electrophysiology. 2016; 21(11):1573–88. Available from: http://dx.doi.org/10.1038/mp.2016.158

32. Malik N, Rao MS. A review of the methods for human iPSC derivation. Methods in Molecular Biology. 2013; 997(5):23–33.

33. Shi Y, Inoue H, Wu JC, Yamanaka S. Induced pluripotent stem cell technology: A decade of progress. Nature Reviews Drug Discovery. 2017; 16(2):115–30.

34. Takahashi K, Yamanaka S. Induction of Pluripotent Stem Cells from Mouse Embryonic and Adult Fibroblast Cultures by Defined Factors. Cell. 2006; 126(4):663–76.

35. Takahashi K, Tanabe K, Ohnuki M, Narita M, Ichisaka T, Tomoda K, et al. Induction of Pluripotent Stem Cells from Adult Human Fibroblasts by Defined Factors. Cell. 2007; 131(5):861–72.

36. Trujillo CA, Muotri AR. Brain Organoids and the Study of Neurodevelopment. Trends in Molecular Medicine [Internet]. 2018; 24(12):982–90. Available from: https://doi.org/10.1016/j.molmed.2018.09.005

37. Velasco S, Kedaigle AJ, Simmons SK, Nash A, Rocha M, Quadrato G, et al. Individual brain organoids reproducibly form cell diversity of the human cerebral cortex. Nature [Internet]. 2019;. Available from: http://www.nature.com/articles/s41586-019-1289-x

38. Nogueira GO, Garcez PP, Bardy C, Cunningham MO, Sebollela A. Modeling the Human Brain With ex vivo Slices and in vitro Organoids for Translational Neuroscience. Frontiers in Neuroscience [Internet]. 2022; 16. Available from: https://www.frontiersin.org/articles/10.3389/fnins.2022.838594/full

39. Flecknell P. Replacement, reduction and refinement. ALTEX [Internet]. 2002; 19(2):73–8. Available from: http://www.ncbi.nlm.nih.gov/pubmed/12098013

40. Li M, Jancovski N, Jafar-Nejad P, Burbano LE, Rollo B, Richards K, et al. Antisense oligonucleotide therapy reduces seizures and extends life span in an SCN2A gain-of-function epilepsy model. Journal of Clinical Investigation [Internet]. 2021; 131(23). Available from: https://www.jci.org/articles/view/152079

41. Schoch KM, Miller TM. Antisense Oligonucleotides: Translation from Mouse Models to Human Neurodegenerative Diseases. Vol. 94, Neuron. Cell Press; 2017. p. 1056–70.

42. Papasaikas P, Valcárcel J. The Spliceosome: The Ultimate RNA Chaperone and Sculptor. Vol. 41, Trends in Biochemical Sciences. Elsevier Ltd; 2016. p. 33–45.

43. NIHR. AVXS-101 for spinal muscular atrophy. NIHR Innovation Oservatory Evidence Briefing. 2018; (Aprol):1–8.

44. FDA. FDA approves innovative gene therapy to treat pediatric patients with spinal muscular atrophy, a rare disease and leading genetic cause of infant mortality.

45. Tabrizi SJ, Leavitt BR, Landwehrmeyer GB, Wild EJ, Saft C, Barker RA, et al. Targeting Huntingtin Expression in Patients with Huntington’s Disease. New England Journal of Medicine. 2019; 380(24):2307–16.

46. Kingwell K. Double setback for ASO trials in Huntington disease. Vol. 20, Nature reviews. Drug discovery. NLM (Medline); 2021. p. 412–3.

